# *Klp61f* and *ncd* function as an accelerator and brake to regulate myonuclear spacing

**DOI:** 10.64898/2026.07.24.740572

**Authors:** Jorel R. Padilla, Yunshu Qiu, Gretchen Kimmel, Mackenzie Vallely, Lydia Olivieri, Eric S. Folker

## Abstract

One of the first genes identified to regulate the spacing of nuclei in the multinucleated myofiber was *Kinesin-1*. However, the mechanism by which *Kinesin-1* or other kinesins regulate myonuclear spacing is not known. Critically, the myofiber lacks centrosomes, and the many myonuclei act as the primary microtubule organizing centers of the cell. Because of this unique re-structuring, we hypothesized that the kinesins that drive centrosomes apart during mitotic spindle elongation may play a similar role in spacing myonuclei. We found that the bipolar *Kinesin-5* (*Klp61f*) and the (-)-end directed *Kinesin-14* (*ncd*) were both necessary for myonuclear spacing at different times, with both being necessary during embryogenesis, but only *ncd* being necessary in the fully differentiated myofiber. To investigate the shared mechanisms during embryogenesis, we used live-imaging and found that, similar to the mitotic spindle, *Klp61f* acts as an accelerator for myonuclear movement, whereas *ncd* acts as a brake contrary to this movement. To investigate these mechanisms and test the hypothesis that this is dependent on microtubule-sliding, we used super-resolution microscopy to visualize and quantify the microtubule network in animals with disrupted *Klp61f* or *ncd*. We found that in both cases, there was a decrease in the amount of microtubule overlap between neighboring myonuclei. Furthermore, we found that disrupting *ncd* led to severe changes in microtubule network organization, supporting our hypotheses that microtubule-sliding is necessary to space myonuclei, and that *ncd* likely functions through a unique mechanism in the differentiated myofiber to maintain myonuclear spacing. Together, our data supports a model where myonuclear spacing is regulated by a counteracting force generated by different kinesins during embryonic development. Furthermore, one kinesin, *ncd*, is repurposed in the differentiated myofiber to dynamically crosslink microtubules, a function necessary to anchor nuclei in place. Thus, kinesin motors regulate myonuclear spacing across developmental time by leveraging opposing forces through diverse mechanisms.

## Introduction

The movement and positioning of the nucleus are conserved processes observed in all eukaryotes (Gundersen and Worman, 2013). This is of particular interest in the multinucleated myofiber, which has multiple nuclei evenly spaced throughout the myofiber. This spacing is established during embryogenesis and maintained in terminally differentiated myofibers, and is crucial to myofiber function (Bruusgaard et al., 2003). Critically, a hallmark of many muscle diseases is the loss of the even nuclear spacing observed in the myofiber (Puckelwartz et al., 2009). Outside of the context of disease, the evenness of myonuclear spacing is decreased in aged and damaged muscle (Buckley et al., 2022; Cristea et al., 2010). Together, these examples illustrate a strong correlation between the evenness of myonuclear spacing and the function of the myofiber. Therefore, elucidating the mechanisms that move and space nuclei is crucial to understanding the development of the myofiber as well as the onset and progression of disease.

Mechanistically, this even spacing is conserved from mammals to *Drosophila* and is dependent on the Linker of Nucleoskeleton and Cytoskeleton (LINC) complex (Cadot et al., 2012; Gimpel et al., 2017; Wang et al., 2015; Wilson and Holzbaur, 2015). The LINC complex spans the nuclear envelope (NE), simultaneously binding to lamins and chromatin in the nucleoplasm and the cytoskeleton in the cytoplasm, providing the means to transmit force between the cytoplasm and the nucleus. Consistent with the hypothesis that the even spacing of nuclei is critical to myofiber health, disruptions in genes that encode the LINC complex have been linked to multiple muscle diseases and dystrophies and animal models of these diseases show disrupted nuclear spacing (Collins and Mandigo et al., 2017; Folker and Baylies, 2013; Puckelwartz et al., 2009; Stroud et al., 2017; Zhang et al., 2007).

In addition to the LINC complex, myonuclear spacing requires microtubules, microtubule associated proteins, and microtubule motors (Cadot et al., 2012; Collins et al., 2021; Folker et al., 2014; Metzger et al., 2012; Rosen et al., 2019; Wilson and Holzbaur, 2012; Wilson and Holzbaur, 2015). Despite the growing list of genes that contribute to myonuclear movement, the mechanisms by which the microtubule network is used by microtubule motors to move myonuclei is not known. Consistent with this, disruption of different genes results in distinct phenotypes. For example, Kinesin-1 was the first motor identified to regulate myonuclear movement (Metzger et al., 2012). However, the phenotype in the *Khc^8^* null animal is weak compared to the phenotype when the expression of either *Ensconsin/MAP7* or *Klarsicht* is disrupted. This suggests that additional kinesin motors may contribute to myonuclear movement.

Although the mechanisms of myonuclear movement are conserved, *Drosophila* provides extensive genetic resources and external development which makes it ideal for testing how nuclei move within the contexts of an actively developing tissue. The active positioning of nuclei happens over distinct steps in development. Either immediately before or during fusion of myoblasts into the growing myotube, the centrosome is broken down and the centrosomal components are then relocalized to the NE, establishing the nucleus as an MTOC (Gimpel et al., 2017; Srsen et al., 2009). After fusion, the newly incorporated myonuclei are moved to the center of the developing myotube by a microtubule-dependent mechanism (Aguilar et al., 2013; Gimpel et al., 2017; Roman and Gomes, 2018). This single cluster of nuclei then separates into two equal clusters that move to the opposite poles of the muscle (Folker et al., 2012; Metzger et al., 2012). The separation of a single cluster into two clusters and the movement towards the muscle ends requires microtubules, the microtubule (+) end-directed motor *Kinesin-1* (*KIF5B*), and the microtubule (-) end-directed motor cytoplasmic *Dynein* (Folker et al., 2014; Metzger et al., 2012).

The poleward movement of myonuclei in the myotube and its dependence on microtubules and motors is reminiscent of the mitotic spindle, in which centrosomes are moved away from each other and toward the cell ends (Mann and Wadsworth, 2019; McIntosh et al., 1969; Östergren et al., 1960). At its core, the elongation of the spindle is characterized by the movement of two microtubule organizing centers (MTOCs) apart and toward the cell cortex by mechanisms that are dependent on several specialized kinesins. Considering that the myonuclei function as MTOCs in myotubes, myonuclear movement could be described in precisely the same way. Therefore, we have investigated whether several mitotic kinesins, kinesins with defined functions in the organization and elongation of the mitotic spindle, also regulate nuclear movement in the postmitotic myotube.

Here we demonstrate that two mitotic kinesins, the (-) end-directed *Kinesin-14* (*ncd* in *Drosophila*) and the (+) end-directed bipolar *Kinesin-5* (*Klp61f* in *Drosophila*) are necessary for myonuclear spacing. Mechanistically, we demonstrate that these motors work in opposition to regulate the speed at which myonuclei are moved during embryonic muscle development. Furthermore, we show that *ncd* is required for microtubule bundling and broader microtubule network organization in the differentiated myofiber. Taken together, our data suggests that the microtubule sliding mechanism described in mitotic spindle elongation is conserved in the post-mitotic myofiber, wherein myonuclear movement and spacing is regulated by several kinesins.

## Results

### *Klp61f* and *ncd* regulate myonuclear spacing at different developmental stages

While we and others have shown that myonuclear spacing is microtubule- and motor-dependent, the mechanisms of myonuclear movement are still poorly understood (Collins et al., 2021; Folker et al., 2014; Levy and Holzbaur, 2008; Metzger et al., 2012; Wilson and Holzbaur, 2012; Wilson and Holzbaur, 2015). For example, although Kinesin-1 regulates several aspects of myonuclear movement (Folker et al., 2014; Wilson and Holzbaur, 2012; Wilson and Holzbaur, 2015), whether there are roles for other kinesins has not been tested. To address this gap, we used RNAi mediated depletion to test whether the mitotic kinesins *Klp61f, Klp3A, Klp10A, ncd,* and *Pavarotti* (Adams et al., 1998; Del Castillo et al., 2015; Ems-McClung et al., 2025; Radford et al., 2017) contribute to myonuclear spacing. All RNAi were previously validated and we provide additional validation by qPCR (Table 1 & Fig S1A) (Davidson et al., 2023; Deshpande and Telley, 2021; Karabasheva and Smyth, 2019; Radford et al., 2012).

**Table 1.**

| Targeted Gene | Specific RNAi | Validation Evidence |
| --- | --- | --- |
| *ncd* | B58144 | This publication, Figure S1 A&B |
| Klp61f | v109280 | Karabasheva & Smyth, 2019,<br>Figure S2 |
| Pav | B42573 | Davidson et al, 2023; Figure S1 |
| Klp3a | B40944 | Deshpande et al, 2021; Table 1 |
| Klp10a | B33963 | Radford et al, 2012; Figure 2B |

We first used Twist-Gal4 to express each UAS-RNAi in the mesoderm when there is active translocation of myonuclei to determine which kinesins are necessary for embryonic nuclear movement. We then measured the spacing of myonuclei in differentiated larval myofibers as previously described (Collins & Mandigo et al., 2017; Mandigo et al., 2019; Padilla et al., 2025). Expression of RNAi against *ncd* or *Klp61f* during active myonuclear movement disrupted the evenness of myonuclear spacing in differentiated larval myofibers (Figs. 1 A,C), but the expression of RNAi against each of the other kinesins had no effect on myonuclear spacing. We interpret this to mean that *Klp61f* and *ncd* are necessary for the embryonic movement of myonuclei during myofiber development.

**Figure 1:**
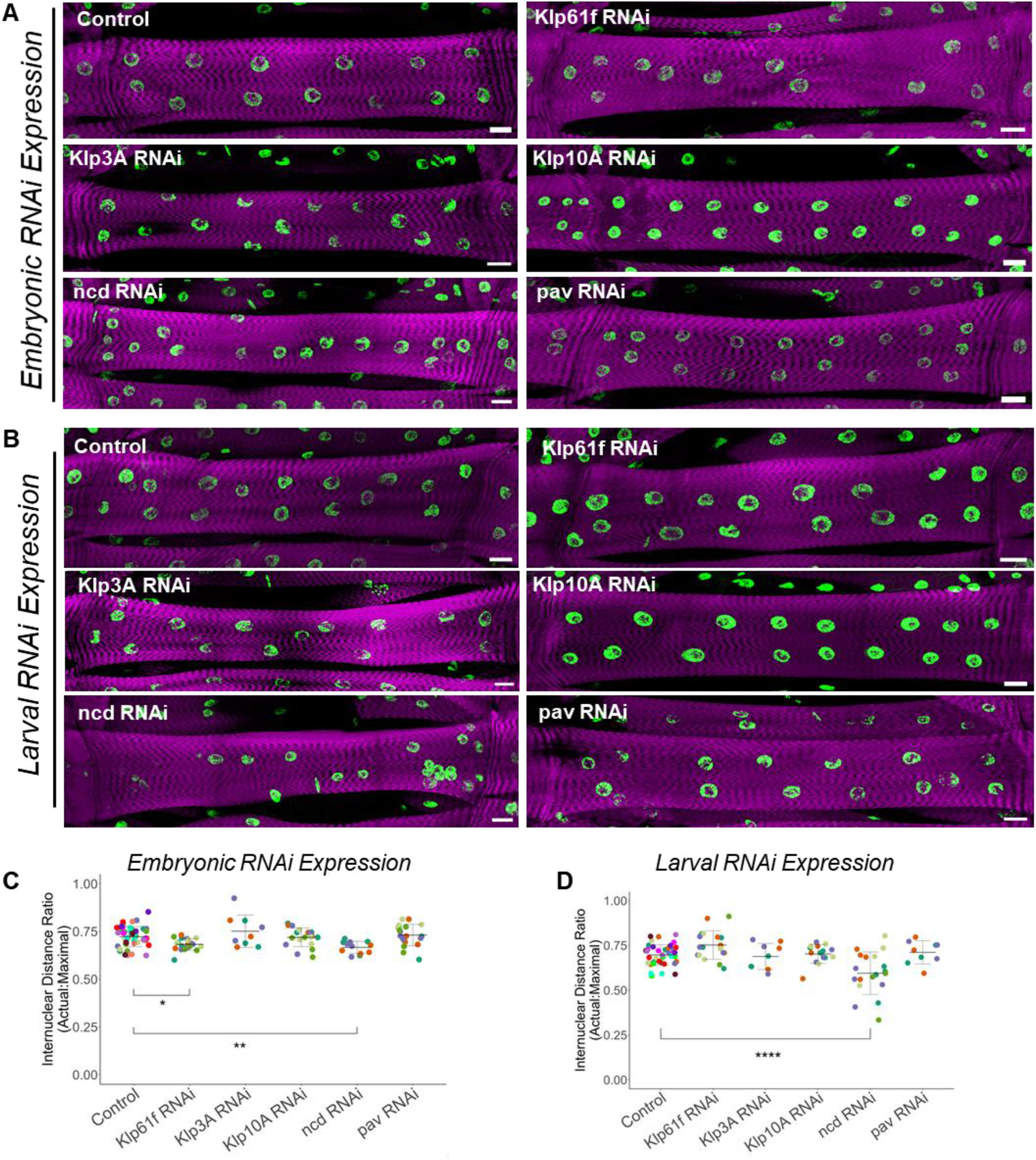
*Kinesin-5* and *ncd* are both essential to achieve myonuclear spacing, but only *ncd* is necessary to maintain myonuclear spacing. (A) Fluorescence images of VL3 muscles from L3 larvae expressing RNAi against indicated gene under control of the embryonic mesodermal-specific driver Twist-GAL4. Magenta, phalloidin/muscles; green, Hoechst/nuclei. Scale bar = 25μm **(B)** Fluorescence images of VL3 muscles from L3 larvae expressing RNAi against indicated gene under control of the larval muscle-specific driver Mhc-GAL4. Magenta, phalloidin/muscles; green, Hoechst/nuclei. Scale bar = 25μm **(C)** Internuclear distance ratio in larval muscles expressing the indicated RNAi under control of the embryonic mesodermal-specific driver Twist-GAL4. Each point is one myofiber. Myofibers from the same animal are the same shape and color. At least three myofibers were analyzed from a minimum of 3 L3 larvae. Data compared by ANOVA. * = p < 0.05. **(D)** Internuclear distance ratio in larval muscles expressing the indicated RNAi under control of the larval muscle-specific driver Mhc-GAL4. Each point is one myofiber. Myofibers from the same animal are the same shape and color. At least three myofibers were analyzed from a minimum of 3 L3 larvae. Data compared by ANOVA *** = p < 0.001.

We have shown that myonuclear movement and the maintenance of myonuclear spacing are mechanistically distinct (Padilla et al., 2025). Therefore, we tested whether any kinesins were required to maintain myonuclear spacing in the differentiated myofiber. To test this, we used Mhc-GAL4 to drive the expression of the same RNAi constructs during the larval stages. Expression of RNAi against *ncd* caused a significant disruption to nuclear spacing, but the other kinesins, including *Klp61f*, were not necessary to maintain myonuclear spacing during larval muscle development (Fig. 1 B,D). Taken together, these data show that while both *Klp61f* and *ncd* are both necessary to achieve myonuclear spacing, only *ncd* is necessary for maintaining myonuclear spacing.

### Myonuclear spacing in the myotube is dependent on both *ncd* and *Klp61f*

Because disrupting either *ncd* or *Klp61f* during embryogenesis resulted in poorly spaced myonuclei, we next examined the developing myotubes in stage 16 embryos. We measured the distance between the myonuclear clusters and the muscle ends as previously described (Auld et al., 2018; Collins et al., 2021; Padilla et al., 2025) and found that animals with disrupted *ncd* expression had myonuclear clusters that were closer to both the ventral and dorsal ends of the muscle (Figs. 2 A,B,C). Interestingly, disruption of *Klp61f* led to no change in the distance between myonuclear clusters and the ends of muscles. This was unexpected as disruption of *Klp61f* during embryogenesis showed a difference in myonuclear spacing in the differentiated myofiber (Figs. 1 A,C). However, our analysis is limited in that we only measured nuclear position at a specific developmental time. Additionally, the size of the myotube may not allow for detection of smaller, yet biologically significant changes in myonuclear position. Because we see a clear phenotype in the larval myofiber, it is likely that the effect of disrupting *Klp61f* is not observable by this specific assay in stage 16 myotubes.

**Figure 2:**
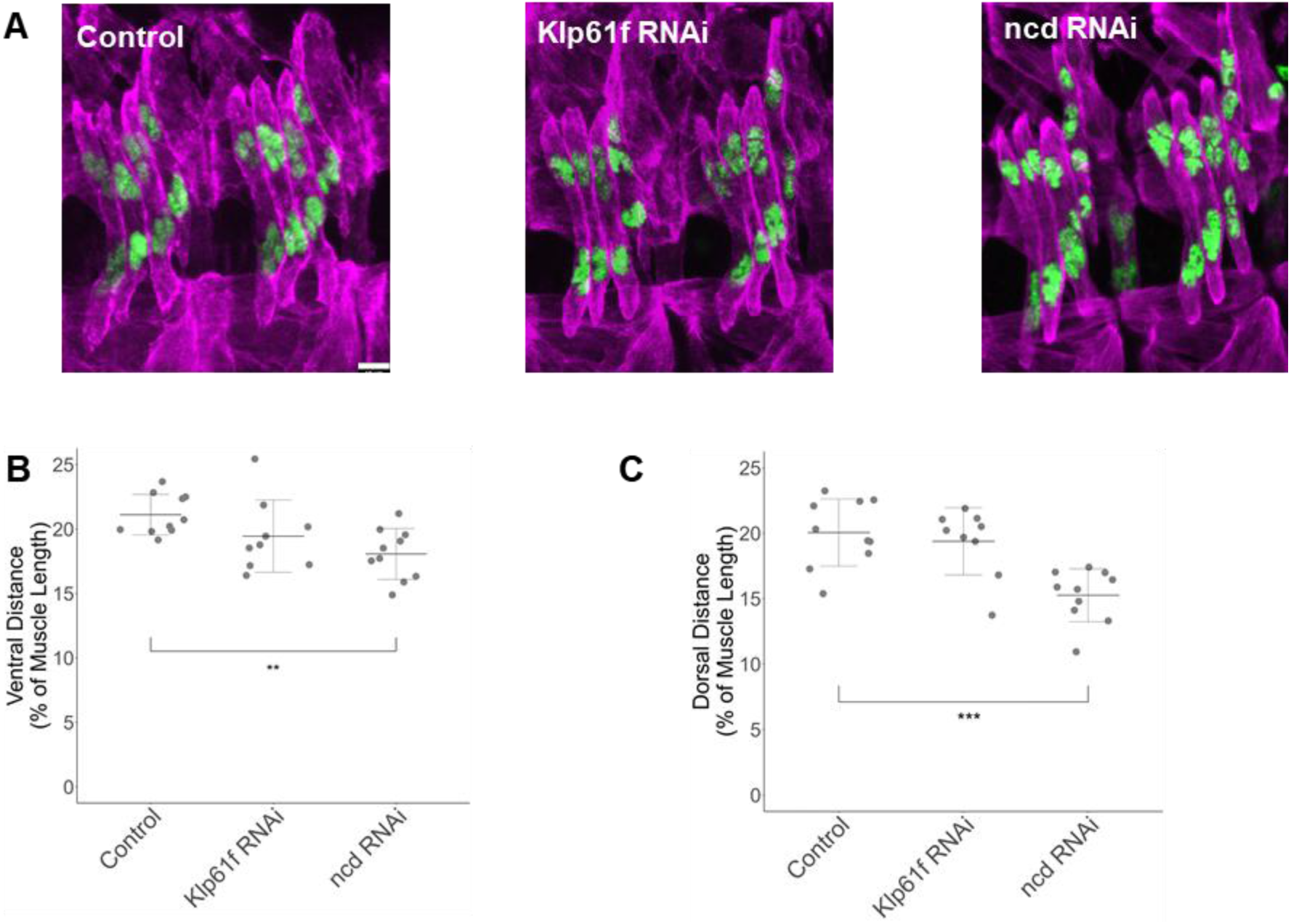
*ncd* but not *Klp61f* regulates nuclear position in embryos. **(A)** Immunofluorescence images of LT myotubes in stage 16 embryos expressing the indicated RNAi under control of the embryonic mesodermal-specific driver Twist GAL4. Magenta, tropomyosin/muscles; green, apRed/nuclei. Scale bar = 10μm. **(B)** Average distance from the ventral end of the myotube to the nearest nuclear cluster. Data compared by ANOVA. ** = p < 0.01 **(C)** Average distance from the dorsal end of the myotube to the nearest nuclear cluster. Data compared by ANOVA. *** = p < 0.001 **B & C**: Each point is the average of LT1, LT2, LT3 from four hemisegments in a single embryo. RNAi is under the control of the embryonic mesodermal-specific driver Twist-GAL4. At least 7 animals were analyzed per genotype.

### *Klp61f* and *ncd* act as a dynamic accelerator and brake respectively to regulate the separation of myonuclei

The measurements of the distance between myonuclear clusters and myotube poles is a measure of where these myonuclei are at a fixed endpoint. To better understand how *Klp61f* and *ncd* contribute to myonuclear spacing, we measured the speed at which myonuclear clusters separate in developing embryos when either *Klp61f* or *ncd* expression is disrupted. In animals with disrupted *Klp61f* expression, the clusters separated more slowly than in controls (Fig. 3 A,B), suggesting that Klp61f produces a force that drives clusters apart. Conversely, in animals with disrupted *ncd* expression, myonuclear clusters separated more rapidly than in controls (Fig. 3 A,B), suggesting that ncd produces a force that restricts the separation of clusters. Because both *Klp61f* and *ncd* regulate the speed of myonuclear cluster separation, we asked if this translated to differences in the distance traveled by individual myonuclei. To answer this question, we measured the total distance traveled by individual myonuclei over 1.5 hours. We found that in animals with disrupted *Klp61f*, individual myonuclei traveled a shorter distance than controls (Fig. 3C). In animals with disrupted *ncd*, individual myonuclei traveled a greater distance than controls (Fig. 3C). Together these data suggest that Klp61f acts as an accelerator to move individual myonuclei away from one another whereas ncd acts as a brake to restrict that movement. This allows for dynamic regulation of myonuclear spacing by the regulation of opposing forces.

**Figure 3:**
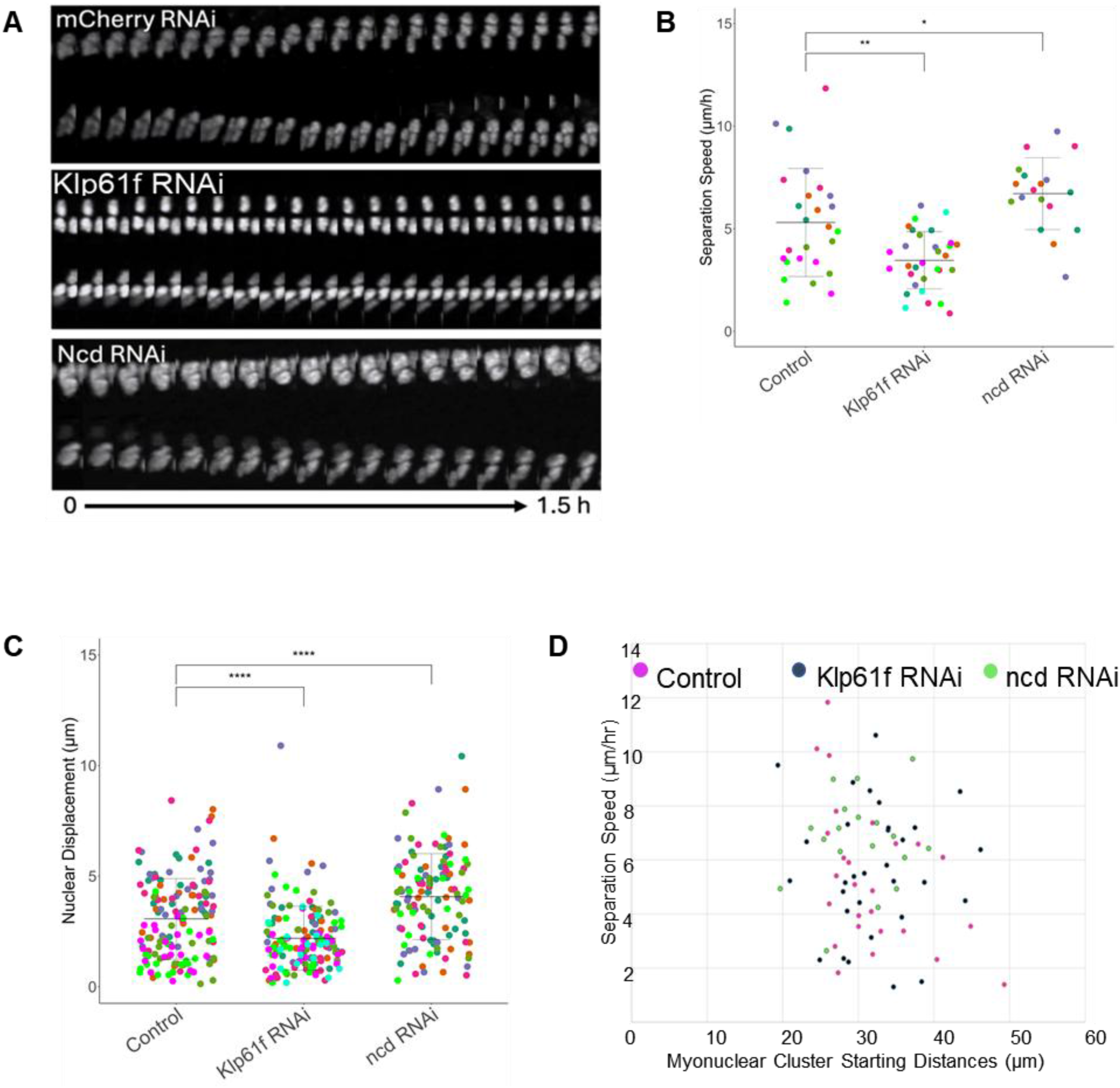
Nuclear separation speed is tightly regulated by both minus- and plus-end directed kinesins. **(A)** Montages of myonuclear clusters separating in stage 16 embryos expressing RNAi against indicated gene under control of the embryonic mesodermal-specific driver Twist-GAL4. White; NLS-DsRed **(B)** Average separation speeds of myonuclear clusters in the indicated genotypes. Each point is one measurement taken from one LT muscle. Data compared by ANOVA. * = p < 0.05; ** = p < 0.01 **(C)** Graph showing the average displacement of individual nuclei after a 90-minute acquisition in the indicated genotypes. Each point is one nucleus. Points of the same color are from the same LT muscle. RNAi expression is under the control of by Twist-GAL4. Data compared by ANOVA. **** = p < 0.0001. **(D)** Graph showing the relationship between nuclear cluster speed and the initial starting distances of nuclear clusters in the indicated genotypes. Each point is one nucleus in an LT muscle from a stage 16 embryo.

Importantly, we considered that these data may be due to differences in the initial distance between myonuclei at the onset of the measurements. As myonuclei move away from one another, the additional space could allow for an increase in the total length of microtubule overlap between myonuclei. The increased overlap of microtubules could allow for increased binding of motors and greater force production. We measured the speed of the separation of myonuclear clusters as a function of the initial distance between clusters and found no correlation between movement speed and the initial distance in controls or in embryos with disrupted *Klp61f* or *ncd* expression (Fig. 3D). Taken together, our data indicates that Klp61f and ncd act as an accelerator a brake respectively to regulate the speed at which myonuclear clusters separate. Furthermore, the balance of these activities is critical for establishing the even spacing of myonuclei in the differentiated myofiber as shown in Figures 1 and 2.

### *Klp61f* and *ncd* differentially regulate the microtubule network organization necessary for microtubule-sliding

While we identified *Klp61f* and *ncd* as novel regulators of myonuclear movement, the mechanism remained unclear. Microtubule sliding is essential for force production in several tissues and cell types (Del Castillo et al., 2015; Leary et al., 2019; Lu et al., 2018; Weinger et al., 2011; Winding et al., 2016). This is most notable and best described in the mitotic spindle (Ems-McClung et al., 2025; Leary et al., 2019; Liu et al., 2021). During mitosis, Klp61f and ncd first bind antiparallel microtubules simultaneously and then produce a net force which moves the microtubules in opposite directions, a process termed microtubule sliding (Liu et al., 2021; Radford et al., 2017). However, although microtubule sliding has been proposed as a mechanism for myonuclear spacing (Metzger et al., 2012), microtubule sliding has never been demonstrated in the myotube or the myofiber. A prerequisite for microtubule sliding is the presence of overlapping antiparallel microtubules. We used AiryScan microscopy to visualize microtubule organization between adjacent myonuclei and examine whether there were incidents of overlapping microtubules. We traced individual microtubules from each nucleus and found many examples where microtubules from neighboring nuclei overlapped (Figs. 4 A,A’,A’’). When we expressed RNAi against either *Klp61f* or *ncd*, the frequency of microtubule overlaps between myonuclei decreased (Fig. 4B). To confirm the presence of microtubule overlaps, we measured the width of microtubules as a function of distance. In control animals, we found that at the points of overlap, the microtubules doubled in thickness, implying an antiparallel overlap (Figs. 4 A’’,B,C,D). However, in animals with disrupted *ncd* or *Klp61f*, the maximum width was lower due to the presence of fewer overlaps. Furthermore, the length of the overlap was shorter when either *Klp61f* or *ncd* expression was disrupted. Consistent with the increase in the width of the tubulin signal, there is an increase in the intensity of the tubulin signal in the same locations, further indicating the presence of microtubule overlap (Figs. 4 C,D,E). Interestingly, in animals with disrupted *ncd* expression, there were fewer microtubule overlaps than in animals with disrupted *Klp61f* expression (Fig. 4B). This more severe phenotype in animals with disrupted *ncd* may explain why *ncd*, but not *Klp61f*, is necessary for maintaining myonuclear spacing in the fully differentiated myofiber (Figs. 1 B,D & 4 A,B,D,E).

**Figure 4:**
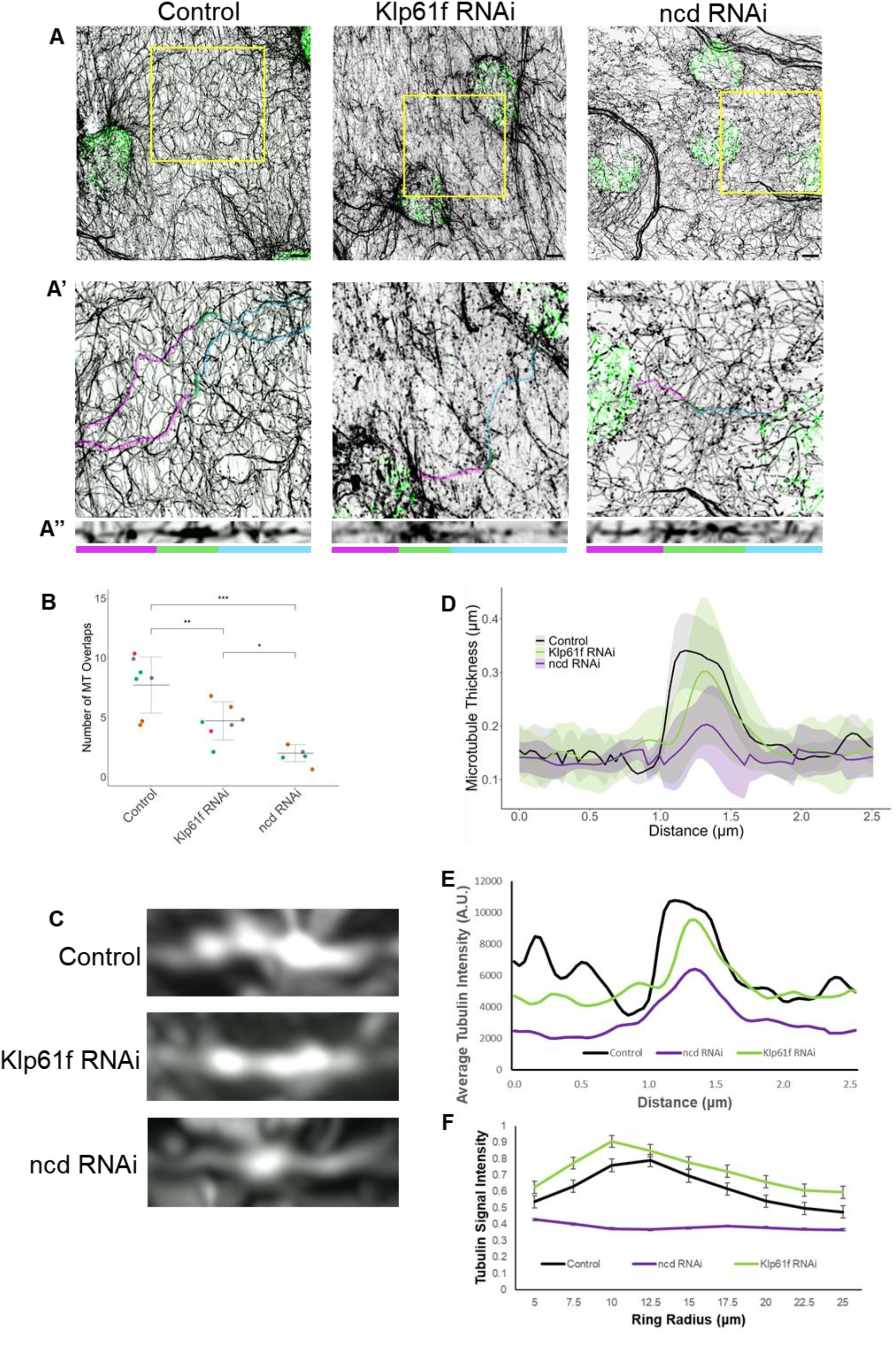
Microtubule network organization is differentially dependent on both *Klp61f* and *ncd.* **(A)** Immunofluorescence AiryScan images of adjacent nuclei from a VL3 muscle in an L3 larvae expressing indicated RNAi under control of the larval muscle-specific driver Mhc-GAL4. Green, Hoechst; black, α-tubulin. Scale bar = 5μm **(A’)** Immunofluorescence AiryScan images of zoomed-in microtubule networks indicated by yellow boxes seen in **A**. Cyan and magenta highlighting traces individual microtubules from adjacent myonuclei. Green highlighting shows areas of microtubule overlap. **(A’’)** Microtubule filaments from **A’** traced and straightened to show microtubule overlap. Color-coding below corresponds to the highlighting in **A’**. **(B)** Graph showing the number of microtubule overlaps between two nuclei. Each point corresponds to the area between two nuclei in one muscle. All points are in different muscles and points of the same color are from the same animal. Data compared by ANOVA. * = p < 0.05; ** = p < 0.01; *** = p < 0.001 **(C)** AiryScan images of microtubule filaments which were traced and then ‘straightened’ depicting the doubling in thickness at points of potential microtubule overlap in VL3 muscles from L3 larvae expressing indicated RNAi under control of the larval muscle-specific driver Mhc-GAL4. White, α-tubulin. **(D)** Graph showing thickness of α-tubulin signal as a function of distance along microtubule filaments in indicated genotypes. **(E)** Graph showing intensity of α-tubulin as a function of distance along microtubule filaments in indicated genotypes. **(F)** Graph showing changes in α-tubulin intensity as a function of distance from the center of the myonucleus in indicated genotypes.

To better understand the effects of disrupting *Klp61f* or *ncd* on the broader organization of the microtubule network, we used AiryScan microscopy to visualize and quantify the organization of the microtubule network beyond counting microtubule overlaps. We and others have previously described a nucleus-associated ring of microtubules (Bruusgaard et al., 2006; Elhanany-Tamir et al., 2012) and linear microtubules extending from this ring (Collins et al., 2021; Padilla et al., 2025) that are crucial for myonuclear spacing. When *ncd* expression was disrupted, there was a decrease in the brightness and thickness of the microtubule ring immediately surrounding the myonucleus (Figs. 4 A,F). Disruption of *Klp61f* did not result in a decrease in the brightness or thickness of the microtubule ring surrounding the myonucleus (Figs. 4 A,F). To better quantify and contextualize these observations, we measured the fluorescence intensity of α-tubulin as a function of distance from the nucleus (Fig. 4F). We found that in animals with disrupted *Klp61f,* microtubule intensity followed a trend similar to that seen in controls, with a strong peak coinciding with the microtubule ring and subsequent decrease of signal when moving away from the myonucleus (Fig. 4F). In animals with disrupted *ncd*, there was a loss of the intensity peak often associated with the myonuclear microtubule ring. Additionally, rather than a steady decay of intensity away from the myonucleus as seen in control animals, the intensity remained flat and consistent, indicating no changes in the microtubule network proximal and distal to the myonucleus (Fig. 4F). Taken together, our data suggests that while both *Klp61f* and *ncd* regulate the number of microtubule overlaps, *ncd* appears to play a larger role in the broad organization of the microtubule network in the differentiated myofiber.

## Discussion

Here, we have shown that two kinesins with well-described functions in the mitotic spindle also regulate cytoskeletal and organelle organization in the post-mitotic myofiber. We show that the bipolar kinesin *Klp61f,* and the (-) end-directed kinesin *ncd,* work in opposition to one another to regulate the speed of myonuclear movement during embryogenesis. Critically, both the accelerator and the brake are required for proper myonuclear spacing days later in the differentiated myofibers of a moving larva, suggesting that myonuclear spacing in the differentiated myofiber relies upon the spacing established days earlier in the embryo. Furthermore, *ncd* is also necessary for maintaining the spacing of nuclei in the fully differentiated myofiber. Beyond myonuclear spacing, we have shown that microtubule overlaps necessary for a microtubule-sliding decrease when disrupting either *Klp61f* or *ncd*. Furthermore, we have shown that *ncd* is a critical regulator of microtubule network organization in the myofiber.

Because *ncd* is critical for microtubule organization, *ncd* may maintain myonuclear spacing through the mechanism we recently described as being dependent on the crosslinking of microtubules and actin by *Dystrophin* and *Msp300* (Padilla et al., 2025). This is consistent with previous work showing that ncd has two separate microtubule binding domains in its tail with one being better suited to slide antiparallel microtubules and the other being ideal to bundle parallel microtubules. (Ems-McClung et al., 2025). It seems likely that the sliding domain is utilized during embryogenesis to regulate myonuclear movement and the bundling domain is used in the differentiated myofiber to build stronger bundles of microtubules to anchor nuclei in position.

Mechanistically, our data is consistent with microtubule sliding contributing to myonuclear spacing. The focus of this work is on *Klp61f* and *ncd*, two specialized and competing motors that have been shown to specifically regulate spindle elongation through microtubule sliding (Mann and Wadsworth, 2019; Radford et al., 2017). *Klp61f* is a bipolar motor with high affinity for overlapping antiparallel microtubules and often described as the driving force of microtubule sliding in the mitotic spindle (Kashina et al., 1996). Additionally, *ncd* has been shown to be critical for regulating mitotic spindle elongation to prevent spindle collapse or collision with other structures (Cullen and Ohkura, 2001; Radford et al., 2017). We have identified and quantified changes in microtubule overlap as a result of disrupting either *Klp61f* or *ncd*, supporting the idea that these motors are necessary for establishing the microtubule architecture necessary to support a microtubule-sliding mechanism. Altogether, the network is organized in a manner to support microtubule sliding and motors with defined roles in microtubule sliding are necessary for myonuclear spacing.

Additional questions and work remain, such as whether other kinesins are necessary for spacing myonuclei at distinct developmental stages. A broader screen of the kinesins would resolve this question. Furthermore, structure-function analysis of *ncd*, *Klp61f*, and any additionally identified regulators of myonuclear spacing would provide more definitive mechanistic insight regarding each individual kinesin motor. Finally, coupling these approaches with advanced live imaging could leverage photo-convertible fluorophores to visualize and measure specific changes in microtubule organization and movement to develop a comprehensive understanding of the mechanisms that drive changes in the microtubule network, move myonuclei, and how these two separate behaviors are linked.

## Materials and Methods

### Drosophila genetics

All stocks were grown under standard condition at 25°C. Stocks used were *apRed* which expresses *DsRed* fused to a nuclear localization signal downstream of the *apterous* mesodermal enhancer allowing for visualization of nuclei within the LT muscles during embryonic stages (Richardson et al., 2007), UAS-mCherry RNAi (35785; Bloomington *Drosophila* Stock Center), UAS-ncd RNAi (B58144; Bloomington *Drosophila* Stock Center), UAS-Klp61f RNAi (v109280; Vienna *Drosophila* Stock Center), UAS-Klp3a RNAi (B40944; Bloomington *Drosophila* Stock Center), and UAS-Pav RNAi (B42573; Bloomington *Drosophila* Stock Center). All UAS expression was driven specifically in larval muscle using *Mhc-GAL4*. Controls were *twist-GAL4*, *apRed* or *Mhc-GAL4,* driving *UAS-mCherry RNAi* as indicated in figure legends. The *twist-GAL4*, *apRed Drosophila* line was made by recombining the *apRed* transgene and the specific GAL4 driver. *Mhc-GAL4* is located on the third chromosome. Tubulin Gal4 was used to drive RNAi for qPCR analysis.

### Sample Preparation & Immunohistochemistry

Embryos were collected at 25°C and then dechorionated by submersion in 50% bleach for 5 min. Embryos were then washed with water and then fixed in 10% formalin (HT501128; Sigma-Aldrich) diluted 1:1 with heptane and placed on an orbital shaker that rotated at a rate of 300 rpm for 20 min. Embryos were devitellinized by vortexing in a 1:1 methanol:heptane solution. Embryos were stored in methanol at −20°C until immunostaining. Larvae were dissected as previously described (Mandigo et al., 2019)

Antibodies for embryo staining were used at the following final dilutions: rabbit anti-dsRed, 1:400 (632496; Clontech); rat anti-tropomyosin, 1:200 (ab50567; Abcam). Antibodies for larval staining were used at the following final dilutions: Mouse anti-αTubulin, 1:200 (T6199, Sigma-Aldrich). Conjugated fluorescent secondary antibodies used for embryo and larval staining were Alexa Fluor 488 donkey anti-mouse-IgG (1:200), and Alexa Fluor 647 donkey anti-rabbit-IgG (1:200). Acti-stain 555-conjugated phalloidin (1:400; PHDH1-A; Cytoskeleton) and Hoechst 33342 (1 μg/ml; H3570; Life Technologies) were used for larval staining only. Embryos and larvae were mounted in ProLong Gold (P36930; Life Technologies) and imaged with an Apochromat 40×/1.3 numerical aperture (NA) objective with a 1.0× optical zoom for all embryo and larval confocal images on a Zeiss 880 LSM. Additionally, larval images were acquired with bidirectional 1×3 tile scan with an online stitching threshold of 0.70.

### Analysis of myonuclear position in larvae

We measured myonuclear position in larvae with as previously described (Auld et al., 2018; Collins & Mandigo et al., 2017). First, the area and length of the muscle were measured. Next, the position and number of nuclei were calculated using the multipoint tool in FIJI to place a point in the center of each nucleus. The position of each nucleus was used to calculate the actual internuclear distance. The maximal internuclear distance was then determined by taking the square root of the muscle area divided by the nuclear number. Nuclear number values were retained and graphed separately. This value represented the distance between nuclei if their internuclear distance was fully maximized. The ratio between the actual internuclear distance and the maximal internuclear distance was then used to determine how evenly nuclei were distributed. This method normalized the internuclear distance to both the nuclear count and the muscle area, which resulted in a more representative means of comparison between muscles, larvae, and genotypes. Ventral lateral 3 (VL3) muscles were measured from at least three larvae with at least three VL3 muscles measured from each larva from at least two biological replicates. Statistical analysis was performed with R studio. Student’s *t*-test or ANOVA with multiple comparisons with Dunnett’s post hoc was used depending on number of genotypes.

### Analysis of nuclear position in embryos

The position of nuclei was measured in stage 16 embryos, which is the latest possible stage before cuticle development blocks the ability to perform immunofluorescence microscopy. Embryos were staged primarily by gut morphology as previously described (Folker et al., 2012). Images were processed as maximum intensity projections of confocal *z*-stacks using FIJI. The positioning of the nuclei was measured using the segmented line function in FIJI to determine the distance between either the dorsal end of the muscle and the nearest nucleus or the ventral end of the muscle and the nearest nucleus. LT muscles were measured in four hemisegments from each embryo. At least 7 embryos from each of two independent experiments were measured for each genotype. Statistical analysis was performed with R studio. One-way ANOVA with Dunnett’s post hoc test was used to assess the statistical significance of differences in measurements between experimental genotypes and controls.

### Live Imaging

Embryos for live-imaging were prepared as previously described (Collins et al., 2021). In brief, embryos were collected at 25°C, washed in 50% bleach to remove the outer membrane, washed with water, and mounted with halocarbon oil (Sigma; Product #H8898). For time-lapse imaging of nuclear movement, stage 15 embryos were selected for imaging based on gut morphology, the position of nuclei, and the intensity of the apRed signal as previously described (Auld et al., 2018; Collins et al., 2021; Folker et al., 2012) with the following modifications. Time-lapse images were acquired on a 3i Spinning Disk Confocal Plan-APOCHROMAT 40×, 1.4 NA oil objective with a 1.0× optical zoom at an acquisition rate of 1 min/stack for 1.5 h. Movies were processed in Fiji as maximum-intensity projections of confocal z-stacks and corrected for drift using the Linear stack alignment with the SIFT plug-in. To calculate the separation speed of nuclei, the line tool was used to measure the distance between dorsal and ventral nuclear clusters at time 0 h and again at time 1.5 h. Displacement of each individual nucleus was calculated as the difference between the final and initial positions.

### Airyscan Acquisition & Analysis of Microtubule Organization in Larvae

These three analyses utilized high-resolution images taken using Elyra Airyscan processing. Larval dissections were prepared as previously described and imaged on the same microscope and objective as described above with a 3.0× optical zoom. All step sizes and speed settings were set to ‘optimal’ within the Zen Black acquisition software for every image. Elyra AiryScan post processing was done using Zen Blue.

First, to determine if there was antiparallel microtubule overlap, microtubules from neighboring myonuclei were traced to find regions where the individual filaments met and appeared to double in thickness, implying an overlap. The number of these overlaps was counted.

We further quantified microtubule overlap by tracing microtubules which overlapped with a line of width ‘20’ in FIJI using the segmented line tool followed by the “straighten” function. We then measured the average intensity of microtubule signal as a function of distance along this tracing. At least 5 microtubule overlaps were measured. At least 3 animals were analyzed per genotype.

The second method used to visualize and quantify the organization of the microtubule network aimed at describing changes in the average thickness of microtubule overlap. To achieve this, we created similar linear projections of microtubules as described in our first method. However, we then measured the thickness of the isolated microtubule filament by drawing lines perpendicular to the filament and measuring their length. Filaments were selected randomly in order to reflect the abundance of filaments with overlap in control animals versus the decrease in overlap abundance seen in animals with disrupted genes. Five filaments total from at least three animals were analyzed per genotype.

The third method used to visualize and quantify the organization of the microtubule network used a code which automatically detected nuclei within a myofiber based on Hoechst staining and drew concentric circles of increasing radii from the center of the nucleus. The radius increased by 2.5μm per ring. The mean pixel value within each ring was measured and plotted as a function of circle radius. The mean pixel value corresponds to the fluorescence intensity of α-tubulin. Concentric circle analysis was run on five non-clustered nuclei from three different myofibers across two animals.

### qPCR analysis of RNAi lines

Gene expression was quantified by RT-qPCR as previously described (Mandigo et al., 2019). Briefly, RNA was extracted and isolated from ten L3 larvae by crushing in an Eppendorf tube in 1 ml of TRIzol according to the manufacturer’s instructions (15596026, Invitrogen). A DNase I (04716728001; Sigma-Aldrich) digest was performed on the isolated RNA at 37°C for 30 min according to manufacturer’s instructions. DNase I was inactivated through the addition of EGTA to a final concentration of 2 mM and heating to 75°C for 10 min. RNA integrity and concentrations were determined using the NanoDrop2000 system (Thermo Fisher Scientific). The cDNA library was established by performing reverse transcription using the SuperScript VILO cDNA synthesis kit (11-754-050; Invitrogen), according to the manufacturer’s protocol. The resulting cDNA was used as the template for qPCR using a QuantStudio 3 system (Thermo Fisher) and Power SYBR green PCR Master Mix (4367659, Applied Biosystems) for detection. For each genotype, three technical replicates were performed. Gene transcript levels were quantified using gene-specific primers designed using FlyPrimerBank and primers were validated according to Applied Biosystems’ instructions. The primers used were: *RP49* forward, 5′-GCCCAAGGGTATCGACAACA-3′; *RP49* reverse, 5′- GCGCTTGTTCGATCCGTAAC-3′; *ncd* forward, 5′- ACGAACTGCGTGGTAACAAG- 3′; *ncd* reverse, 5′- CCAGGCGGGATACTTTGCTGC-3′; *Klp61f* forward, 5′- ACACCACCAAGATACGCATTT-3′; *Klp61f* reverse, 5′- CGTAGTGGCTGTTTTGCGAC- 3′; *Pav* forward, 5′- CAGCAGGATGTGTACGCAG-3′; *Pav* reverse, 5′-

GGTGAAAAGCAGACTGTTTCGG-3′; *Klp3a* forward, 5′- AGGATCTATTCGAGACGTGTGT-3′; and *Klp3a* reverse, 5′- GCCCATAGTGTAGGTCTTTCCG-3′. The differences in gene expression were calculated using the ΔΔC_t_ method. Rp49, was used as the reference gene for comparison to the gene of interest for ΔC_t_ values for each sample. Fold changes were expressed as 2^−ΔΔCt^ and plotted in Log_2_ for graphical representation.

---

**Figure S1:**
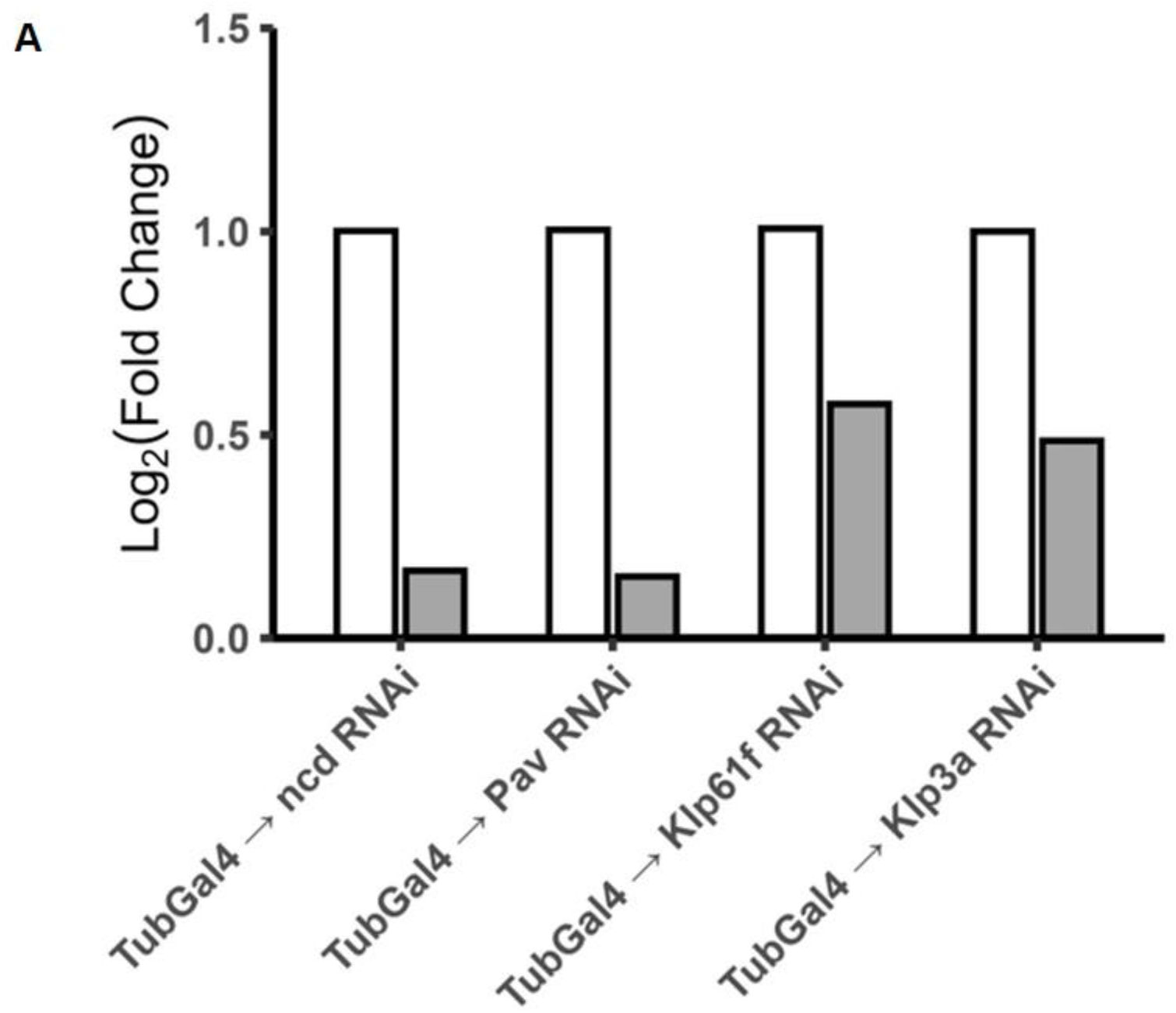
RNAi results in reduction of RNA levels corresponding to genes of interest. **(A)** Fold change in all RNAi lines showing reduction in all target genes. Left white column represents endogenous levels of the gene of interest. The right gray column represents the relative level of gene expression in animals expressing an RNAi against the specified gene. RNAi driven by Tubulin-Gal4.

